# Early life adversity reduces affiliative behavior towards a distressed cagemate and leads to sex-specific alterations in corticosterone responses

**DOI:** 10.1101/2023.07.20.549876

**Authors:** Jocelyn M. Breton, Zoey Cort, Camila Demaestri, Madalyn Critz, Samuel Nevins, Kendall Downend, Dayshalis Ofray, Russell D. Romeo, Kevin G. Bath

**Affiliations:** Columbia University, Department of Psychiatry, New York, NY, USA; Barnard College of Columbia University, Department of Neuroscience and Behavior, New York, NY, USA; Brown University, Department of Cognitive, Linguistic and Psychological Sciences, Providence, RI, USA

**Keywords:** Early life adversity, corticosterone, emotional contagion, social interaction, sex differences

## Abstract

Experiencing early life adversity (ELA) alters stress physiology and increases the risk for developing psychiatric disorders. The social environment can influence dynamics of stress responding and buffer and/or transfer stress across individuals. Yet, the impact of ELA on sensitivity to the stress of others and social behavior following stress is unknown. Here, to test the impact of ELA on social and physiological responses to stress, circulating blood corticosterone (CORT) and social behaviors were assessed in adult male and female mice reared under limited bedding and nesting (LBN) or control conditions. To induce stress, one cagemate of a pair-housed cage underwent a footshock paradigm and was then returned to their unshocked partner. CORT was measured in both mice 20 or 90 minutes after stress exposure, and social behaviors were recorded and analyzed. ELA rearing influenced the CORT response to stress in a sex-specific manner. In males, both control and ELA-reared mice exhibited similar stress transfer to unshocked cagemates and similar CORT dynamics. In contrast, ELA females showed a heightened stress transfer to unshocked cagemates, and sustained elevation of CORT relative to controls, indicating enhanced stress contagion and a failure to terminate the stress response. Behaviorally, ELA females displayed decreased allogrooming and increased investigative behaviors, while ELA males showed reduced huddling. Together, these findings demonstrate that ELA influenced HPA axis dynamics, social stress contagion and social behavior. Further research is needed to unravel the underlying mechanisms and long-term consequences of ELA on stress systems and their impact on behavioral outcomes.

## 1. Introduction

Experiencing early life adversity (ELA) increases the risk for developing psychiatric disorders later in life, including depression and anxiety (Heim and Nemeroff, 2001). One potential mechanism by which ELA can lead to increased risk for these disorders is through changes in stress physiology. In particular, ELA can have a profound impact on the development and functioning of the hypothalamic-pituitary-adrenal (HPA) axis, a key system involved in the body’s response to stress (Koss and Gunnar, 2018; Perry et al., 2019; Sanchez, 2006). The HPA axis plays an essential role in coordinating the physiological stress response, and helps individuals respond appropriately to threatening stimuli. When a threat is detected, a cascade of events occurs that leads to the release of glucocorticoids into the bloodstream; these hormones in turn feedback on this system to terminate the stress response (Herman et al., 2016, 2012). In individuals that have experienced ELA, HPA axis activity, measured via cortisol secretion, is often blunted in response to psychosocial stressors (Bunea et al., 2017; Lovallo, 2013). In addition, ELA can impair the feedback process that typically terminates the stress response (Maccari et al., 2014). However, the effects of ELA on HPA functioning are variable, with evidence supporting both elevated and blunted HPA axis activity (Fogelman and Canli, 2018; Koss and Gunnar, 2018). These opposing findings may be related to differences in approaches to testing this question as well as limited exploration of variables that can impact the stress response, including sex-specific effects (Carpenter et al., 2017; DeSantis et al., 2011) and differences in the social environment. Understanding sex differences in ELA’s effect on the HPA axis is crucial, as males and females exhibit distinct responses to stress (Kudielka and Kirschbaum, 2005) and have variable risk for developing stress-associated pathology (Gobinath et al., 2015; Kessler et al., 2005). Despite this, there is a paucity of studies examining sex-specific effects of ELA on HPA axis functioning.

The individual’s social environment is a critical factor that influences HPA axis functioning. For example, parents and peers can help buffer a stress response, and stress can also be transferred between individuals (Burkett et al., 2016; Hall and Romeo, 2014; Hostinar et al., 2014). Despite the influence of social interactions on the HPA axis, limited research has explored how ELA might interact with the social environment to modulate stress responses. In particular, ELA may influence future sensitivity towards the stress, or emotions, of others, a phenomenon referred to as emotional contagion (Keysers et al., 2022). Few studies have directly examined ELA’s effect on emotional contagion; rather, existing studies have instead focused on ELA’s impact on emotional regulation. For example, children who have experienced ELA are more influenced by emotional contexts, with increased emotional reactivity to negative stimuli (Marusak et al., 2015; Pechtel and Pizzagalli, 2011; Tottenham et al., 2010). However, it remains unknown whether those with a history of ELA might also be more impacted by the stress of another and show a heightened social transfer of stress (emotional contagion), which could in turn impact subsequent behavior, including stress coping strategies. Here too, sex differences have not yet been fully explored, though sex is known to influence stress coping (Tamres et al., 2002).

Animal models of ELA provide a translational tool to test hypotheses and allow us to isolate environmental conditions that may lead to changes in the development and functioning of the HPA axis. Factors such as the social environment as well as the type of adversity experienced can be directly manipulated, with the timing and severity of stress controlled. One common model of ELA, the limited bedding and nesting (LBN) model, leads to fragmented maternal care (Demaestri et al., 2022, 2020; Gallo et al., 2019; Rice et al., 2008), as well as changes in basal corticosterone (CORT) levels in the pups (Bath et al., 2016; Molet et al., 2014; Rice et al., 2008), indicating that HPA axis development is altered following this form of ELA. However, the effects of LBN on CORT levels following a stress challenge have yielded mixed results (Eck et al., 2020; Gilles et al., 1996; McLaughlin et al., 2016). Just as in humans, social factors (such as social or isolated housing during stress exposure) may significantly impact stress responding, thus contributing to this variability. Stress may be experienced both directly by an animal and indirectly, through a social transfer from conspecifics. Yet, there is limited work testing how ELA conditions might influence these two different forms of stress, and how stress experienced in a social context might alter subsequent social behaviors.

In the current study, we sought to understand how ELA in the form of LBN, affects dynamic changes in HPA axis functioning in both a distressed animal directly exposed to footshock stress, and in their unshocked cagemate upon exposure to the distressed cagemate. Further, as social behavior in this context may serve to either buffer or exacerbate the stress of the shocked individual, as well as lead to transfer of stress to the unshocked animal, we also quantified the effects of our manipulation on social behavior following the return of the stressed animal to the cage. To answer these questions, adult male and female mice reared under control or LBN conditions underwent a paradigm where one animal was exposed to a short series of footshocks before being returned to their unshocked cagemate. Blood CORT levels at baseline or at one of two timepoints in both the shocked and unshocked cagemate were then measured to assess dynamic changes in HPA axis response to either footshock stress or to interaction with a shocked cagemate. Lastly, in addition to measuring physiological changes, we also assessed changes in behavioral responding to stress, measuring social behavior between cagemates following this acute stress challenge. Importantly, we also tested for sex differences, helping fill a critical gap in the field. Based on previous literature, we hypothesized that ELA animals would exhibit both a reduction in social interactions following stress as well as increased stress contagion, as measured by CORT levels in the unshocked cagemate. Further, based on prior findings that males and females differentially respond to the LBN paradigm (Bath, 2020; Demaestri et al., 2020; Goodwill et al., 2019, 2018), we expected to observe sex-selective effects of ELA on CORT dynamics following acute stress. By investigating the effects of ELA and sex on HPA axis functioning in a social context, this study contributes to our understanding of key factors that may influence and regulate stress physiology, as well as potential environmental factors that may moderate or exacerbate stress responding. Such work may have important implications for understanding changes in neuroendocrine function associated with risk for psychiatric disorders.

## 2. Methods

### 2.1 Animals and housing

C56BL/6N mice (Charles River) were bred in-house and maintained on a 12h:12h light dark cycle (lights on at 7AM) in a humidity-controlled room, with *ad libitum* access to food and water. Mice were housed in standard Optimice cages (34.3 L x 29.2 W x 15.5 H cm - 484 square cm) with cobb bedding and a 4x4 cm cotton nestlet. Twenty-five litters were included in the current experiments. With the exception of a single litter that had one male pup, all other litters were composed of both male and female pups, ranging from 4 to 12 pups per litter. These litters had a relatively equal mix of male and female animals. All mice were weaned and sex-segregated at postnatal day (p) 21. For the litter with a single male pup, the mouse was paired with an age-matched male mouse from another dam at weaning. 152 adult mice (77 males and 75 females, >p65) were used for the studies described here. A full description of the animals used and their respective dams can be found in Supplemental Table S1. All procedures were approved by the New York State Psychiatric Institute and Columbia University Institutional Animal Care and Use Committees and were in accordance with the National Institutes of Health Guide for the Care and Use of Laboratory Animals.

### 2.2 Limited Bedding and Nesting (LBN)

Male and female breeders were pair-housed in a standard home cage with bedding and a 4x4 cm cotton nestlet. The limited bedding and nesting (LBN) paradigm was conducted as previously described (Bath et al., 2016; Demaestri et al., 2020; Gallo et al., 2019). Briefly, four days following the birth of a litter (p4), the dam and pups were transferred from their standard home cage to a cage with a wire mesh floor, no bedding, and a 3x4 cm cotton nestlet (Figure 1). The wire floor was raised 0.5cm off the bottom of the cage to allow urine and defecations to pass through. Dams and pups remained in these conditions for seven days and on p11 were returned to their standard housing conditions. At weaning, mice were housed in same-sex pairs of either control or LBN conditions until experimental use (>p65). Dams that were assigned to the LBN paradigm remained assigned to LBN for all subsequent litters.

**Figure 1.**
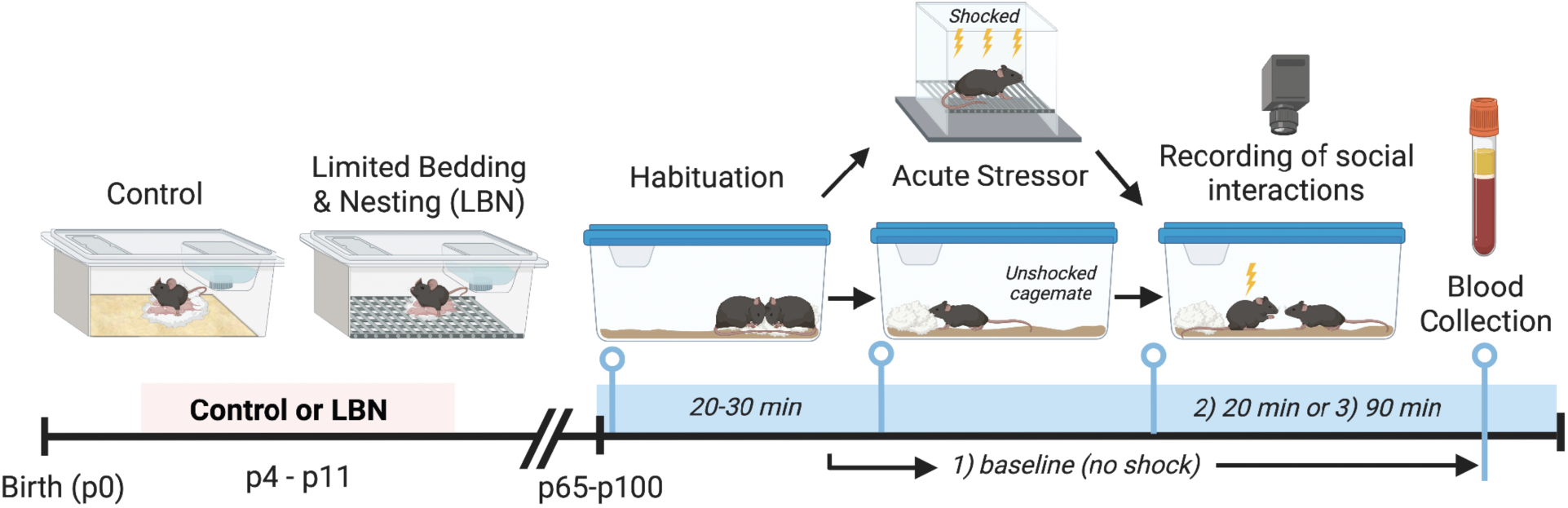
Experimental Timeline. The day of birth was designated as postnatal day (p) 0. On p4, dams and pups assigned to the limited bedding and nesting (LBN) paradigm, were transferred into a new cage with a wire mesh floor and reduced nesting material (a 3x4 cotton nestlet). On p11, dams and pups were returned to normal housing conditions, with standard bedding and a full 4x4cm nestlet. Control litters were left undisturbed during that time. Upon reaching adulthood (>p65), pair housed mice were first habituated to an experimental room. One mouse of the pair then underwent a brief series of footshocks before they were returned to their cagemate. Social interactions were recorded for 20 minutes, and blood was extracted either 20 minutes or 90 minutes following the animals return to their cagemate. For a separate group of mice, blood was collected immediately following habituation as a baseline group.

### 2.3 Experimental Design & Footshock Stress

To test the effects of LBN on stress responses following direct physical stress or following exposure to a stressed mouse, one mouse in a pair-housed cage of control or LBN-reared mice experienced a series of footshocks before returning to their unshocked cagemate. Prior to the footshocks, all animals were removed from the housing room and placed in an adjacent room for a minimum of 20 minutes for habituation. One cagemate was then chosen at random to be ‘Shocked’. This ‘Shocked’ mouse was marked with Sharpie on its tail to distinguish it from its cagemate. The ‘Shocked’ animal was then placed in a Med Associates acoustic startle apparatus equipped with a small animal holder (6.035 x 6.19 x 4.8cm) with grid flooring. Following a one-minute habituation to the chamber, three footshocks (0.5mA) were delivered, with a one-minute inter-trial interval (ITI). Shocked mice were then immediately returned to their home cage with the ‘Unshocked’ cagemate. Blood was collected from both the Shocked and Unshocked cagemate at predetermined timepoints (see serum sampling below) to assess changes in circulating CORT (Figure 1). For a subset of cages (n=37), video recordings of behavior were collected to analyze social interactions (see behavioral scoring below). Shock chambers were cleaned with 70% ethanol between mice.

### 2.4 Serum Sampling

To avoid further stress associated with multiple blood draws, independent groups of mice were used for blood collection at each time point. Blood was collected prior to stress exposure (baseline), at 20 min post stress, or at 90 minutes post stress (n = 5-8 cages per timepoint) from both the Shocked and Unshocked cagemates (Figure 1). All collection procedures occurred between 12:30PM-3:00PM for consistency and to control for circadian effects on CORT levels. Mice from multiple litters were balanced among the baseline, 20- and 90-min groups to control for potential litter effects (Table S1).

To collect blood, animals were first given an intraperitoneal (i.p.) injection of avertin (2,2,2-tribromoethanol, 500mg/kg at 20mg/mL) until non-responsive (approx. one min or less). Trunk blood was then collected following rapid decapitation, within two minutes after injection in order to avoid elevations in CORT due to the injection itself. After collection, blood was allowed to clot for one hour at room temperature and was then centrifuged at 1,000 g for 10 minutes at 4°C. Serum (∼50-150µl) was collected into a clean tube and stored at -20°C for later analysis. The blood collected from two of the mice was not used in the final analysis due to experimental error.

### 2.5 CORT Radioimmunoasssay (RIA)

CORT radioimmunoassays (RIA) were conducted using commercially available kits and reagents and were performed as indicated by the supplier (MP Biomedicals; Solon, OH). All samples were run in duplicate in a single assay, and values were averaged. The lower limit of detectability was 24.03 ng/mL and the intra-assay coefficient of variation was 7.7%. The antibody for CORT provided is highly specific and shows little cross reactivity with other steroid hormones (i.e.,<1%).

### 2.6 Behavioral Scoring

Overhead recordings of homecage behavior were captured either during the habituation period and/or following the acute stressor to examine the social interactions between Shocked mice and their Unshocked cagemates. All videos were processed using Ethovision 16 and were manually scored for social interactions. Both active and passive interactions were scored.

Active interactions consisted of nose-body touching (anytime one mouse’s nose touched any part of the other mouse’s body) as well as touching with the forepaws or mouth (for example, as seen when grooming). Under this umbrella of active interactions, three specific categories were scored: anogenital sniffs, allogrooming and agonistic events. These behaviors are defined as follows:

- Anogenital sniffs were defined as nose to anogenital region contact, initiated by either mouse.
- Allogrooming was defined as grooming of any part of the partner, including the tail, using the mouth and forepaws.
- Agonistic events were defined as wrestling and pinning behavior, as well as biting.

Passive interaction consisted only of huddling behavior, which was defined as follows:

- Huddling (or side-by-side contact) was defined as body-to-body contact with their partner that persisted for longer than 1 second, without engagement in the previously described active interactions.

As the focus was on overall social interaction, these behaviors were scored based on aggregate behavior in each cage and were not based on the behavior of an individual animal. For all animals, approx. 20 minutes of behavior following footshock were analyzed by two independent observers to ensure accuracy. The average inter-rater reliability was >90% concordance in scoring.

### 2.7 Statistical Tests

All data are presented as the mean ± the standard error of the mean (SEM). Outliers were removed using the ROUT method in Graphpad Prism, which combines robust regression with outlier removal. For analysis of CORT, a three-way ANOVA was used to test for effects of rearing condition, sex, and time group separately for shocked and unshocked mice. If a main effect of sex was identified, two-way ANOVAs for rearing condition and time group were conducted separately for males and females. Sidak’s post-hoc analyses were used to correct for multiple comparisons. For behavioral analyses, given known sex differences in social interaction (An et al., 2011; Bredewold and Veenema, 2018), male and female mice were analyzed separately. Thus, unpaired independent sample Welch’s t-tests were used to directly compare behavioral changes between control and footshock stressed animals within each sex. Pearson correlations were used to identify relationships between CORT levels of the two mice within the cage (Shocked vs. Unshocked). In all tests, the alpha value was set at 0.05. All analyses were performed with GraphPad Prism version

8.4 (GraphPad Software, San Diego, California USA) and RStudio (RStudio Team, 2018).

## 3. Results

### 3.1 ELA in the form of LBN led to persistent elevated CORT levels in Shocked females and an increased transfer of stress to their Unshocked cagemate

To examine the physiological response of male and female mice reared under control or ELA conditions to an acute stress exposure, blood was collected from three groups: an unshocked baseline group and two groups following stress exposure: one 20 minutes later and another 90 minutes later. Based on prior literature and preliminary work from our lab, we anticipated to see a peak in CORT levels at 20 minutes and a recovery by 90 minutes (Shanks et al., 1990). Here, exposure to three 0.5mA footshocks produced an increase in CORT over baseline levels at 20 minutes in all Shocked groups (Figure 2A). A 3-way ANOVA yielded a main effect of time group (F_(4,130)_ = 22.77, p<0.0001) and sex (F_(1, 130)_ = 18.49, p<0.0001), and no effect of rearing condition. A sex by rearing condition interaction was also observed (F_(1,130)_ = 6.125, p=0.0146) as was a three-way interaction between time group, sex, and rearing condition (F_(4,130)_ = 3.016, p=0.0204). As a main effect of sex and a sex by rearing condition interaction were observed in our initial analysis, males and females were subsequently analyzed separately.

**Figure 2.**
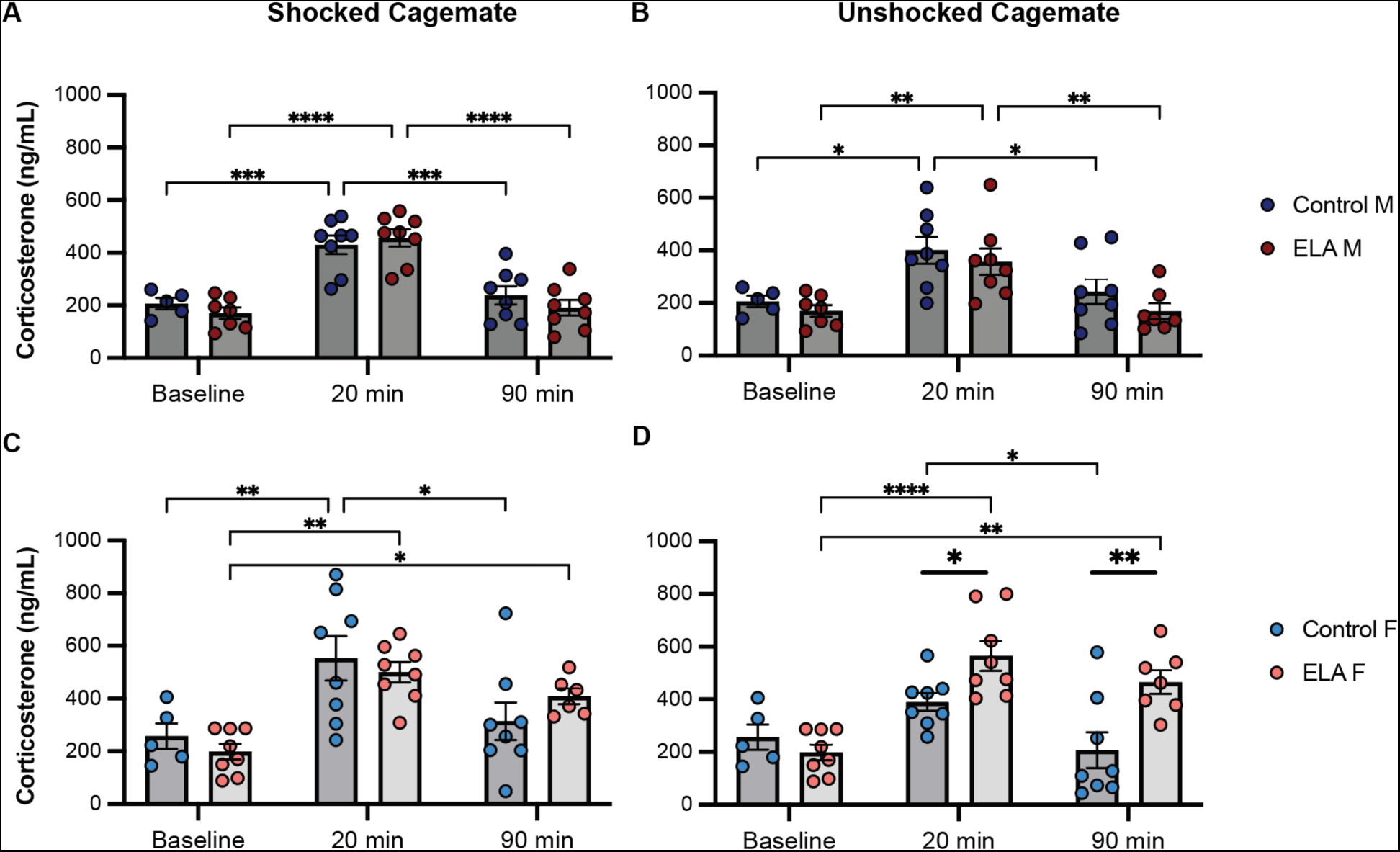
Corticosterone (CORT) levels in Shocked and Unshocked ELA and Control mice. Blood was collected either prior to any stress exposure (baseline) or at 20 min or 90 minutes following footshock stress and circulating corticosterone (CORT) levels were measured in both the Shocked cagemate (A,C) and Unshocked cagemate (B,D) for males and females respectively. A) Shocked ELA and Control males showed a similar increase in CORT at 20 minutes and a return to baseline levels by 90 minutes. B) Unshocked male cagemates in both Control and ELA conditions showed a transfer of stress, with elevated CORT at 20 minutes, and a similar recovery by 90 minutes. C) Shocked ELA and Control females showed a similar increase in CORT at 20 minutes, however at 90 minutes, Control females had returned to baseline levels, while ELA females had not. D) Unshocked ELA females displayed elevated CORT relative to controls at both 20 minutes and 90 minutes, indicating increased emotional contagion and sustained stress responding. Sidak’s post-hoc test: ELA vs. Control. A separate Sidak’s post-hoc test was used to compare CORT at the three different timepoints within each group (bars are shown with smaller lines) (*p<0.05, **p<0.01, ***p<0.001, ****p<0.0001).

Control and ELA-reared males showed similar elevations in CORT levels in response to stress in the form of direct footshock or indirect stress transfer (Shocked: 2-way ANOVA: main effect of time group: F_(2,38)_ = 40.70, p<0.0001; rearing condition: p > 0.05); (Unshocked: 2-way ANOVA: main effect of time group: F_(2,37)_ = 12.77, p<0.0001; rearing condition: p > 0.05); (Figure 2A-B). In both Control and ELA males that were both Shocked and Unshocked, post-hoc tests revealed a statistically significant increase in CORT 20 minutes post stress compared to the baseline groups (Shocked Control: p<0.0001, Shocked ELA: p<0.0001, Unshocked Control: p=0.0129, Unshocked ELA: p=0.008). In addition, CORT levels of the 90-minute group were indistinguishable from the corresponding baseline group for both Shocked and Unshocked mice in both rearing conditions (non-significant, (n.s.), p>0.05). Thus, both Control and ELA-reared male mice showed a significant elevation in CORT in response to stress and a return to baseline by 90 minutes, regardless of the type of stressor (either direct exposure or indirect transfer from their cagemate). Interestingly, the Unshocked cagemates in both Control and ELA conditions showed an identical pattern in the time course of CORT responding, with CORT levels comparable to their Shocked counterparts (3-way ANOVA: n.s. effect of shock condition, p>0.05), indicating a transfer of the stress response in all male mice.

For females, we observed a different pattern of CORT responding as a result of ELA. For mice receiving the shock, we observed a significant effect of time group, but not rearing condition (2-way ANOVA: main effect of time (F_(2,37)_ = 13.20, p<0.0001) n.s. effect of rearing condition, p>0.05; Figure 2C), with similar elevations in CORT at 20 minutes relative to baseline for both Control (p=0.005) and ELA (p=0.0012) mice. By 90 minutes post-shock, CORT levels in the Control females were indistinguishable from the baseline group (n.s., p>0.05); however, Shocked ELA females did not show a similar recovery, and instead remained significantly elevated from baseline levels at 90-minutes (p=0.0486). Further, CORT levels in the 90-minute group were not statistically different from that of the 20-minute group, indicating that Shocked ELA females had not yet recovered to baseline levels after 90 minutes.

Interesting differences in CORT response dynamics were also observed in the Unshocked female groups. A two-way ANOVA revealed a main effect of time group (F_(2,38)_ = 12.47, p<0.0001), a main effect of rearing condition (F_(1,38)_= 9.349, p=0.0041), and a significant interaction between time and rearing (F_(2,38)_ = 4.966, p=0.0121; Figure 2D). Notably, at both 20 and 90 minutes post shock, Unshocked ELA females exhibited greater CORT levels compared to Unshocked Control females (20 min: p = 0.0381, 90 min p = 0.0018), suggesting increased stress contagion in ELA females. In addition, like the Shocked ELA group, Unshocked ELA females did not recover to baseline group levels by 90 minutes post-stress (baseline vs. 90: p=0.0013, 20 vs. 90: n.s., p>0.05). In contrast, CORT levels in Unshocked Control females 90 minutes post-stress were indistinguishable from baseline group levels (baseline vs. 90: n.s., p>0.05). Thus, Unshocked ELA-reared females experience greater stress contagion than controls and show a prolonged elevation in their stress response.

### 3.2 Elevated CORT levels of Shocked mice were predictive of greater CORT in Unshocked cagemates for all groups, except Control females

Circulating levels of CORT provide an index of physiological responding to stress, and the magnitude of the CORT response is thought to correlate with the magnitude of stress experienced. Here, we used a correlation analysis to test whether CORT levels from individual Shocked mice were predictive of CORT levels in their Unshocked cagemate. In this analysis the 20-minute and 90-minute groups were pooled to increase statistical power. Pearson’s correlations revealed a positive correlation between Shocked and Unshocked CORT levels in Control (r = 0.54, p=0.003; Figure 3C) and ELA males (r = 0.65, p=0.0086; Figure 3D), and in ELA females (r=0.59, p=0.027; Figure 3B). Surprisingly, this correlation was not observed in Control females (r = 0.35, p>0.05; Figure 3A). These data suggest that there is a robust transfer of stress between Shocked mice and their Unshocked cagemate for both ELA males and females and for Control males, with the magnitude of CORT responding in Shocked mice positively related to levels observed in Unshocked cagemates.

**Figure 3.**
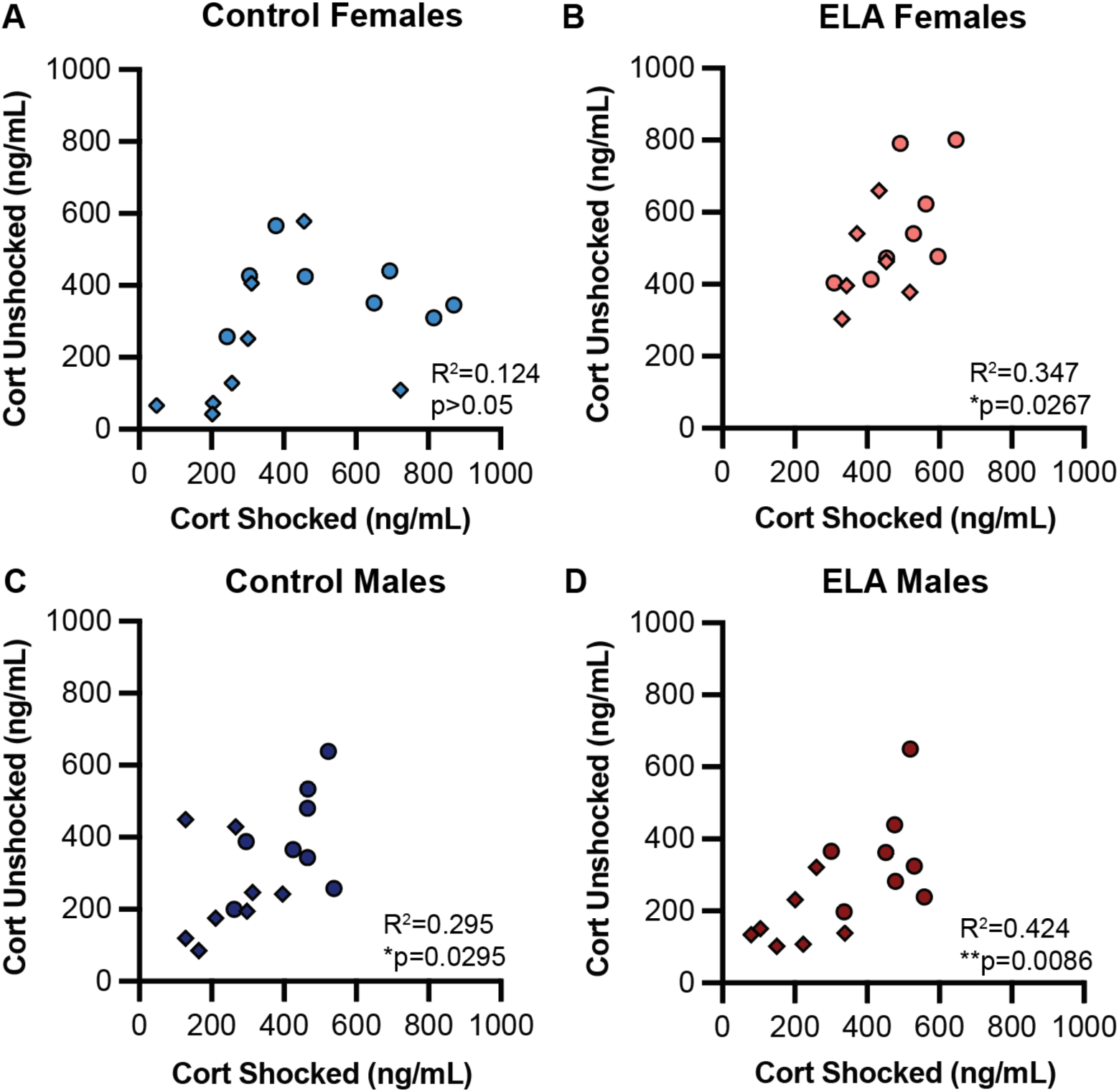
Correlations of CORT between Shocked and Unshocked cagemates. CORT levels between the Shocked and Unshocked mouse in each cage were compared for Control and ELA females (A-B) and Control and ELA males (C-D). A) There was no statistically significant correlation between Shocked and Unshocked CORT levels for control females. For ELA females (B), Control males (C), and ELA males (D) there was a positive correlation between the Shocked mouse’s CORT levels and that of their Unshocked cagemate. (20-minute and 90-minute groups are combined here. Circles indicate animals whose blood was collected at 20 minutes. Diamonds indicate animals whose blood was collected at 90 minutes). (*p<0.05, **p<0.01).

### 3.3 ELA reduced prosocial interactions amongst cagemates following an acute stressor

There is ample work showing that social interactions can influence the stress response; the social environment can be both a source of stress or act to buffer stress (Beery and Kaufer, 2015). Here, we tested for differences in cagemate social interactions following stress exposure across rearing conditions (Control vs. ELA) to gain insight into behavioral outcomes that might explain our observed differences in CORT levels. Both active interactions (including investigative behavior such as nose to body touches, anogenital sniffs, prosocial allogrooming, and fighting), as well as passive side by side huddling were measured. Behavior was continuously scored for 20 minutes upon reuniting the Shocked mouse with its Unshocked cagemate. Given known sex differences in social interaction (An et al., 2011; Bredewold and Veenema, 2018) and the sex differences in CORT levels observed here, male and female mice were analyzed separately.

Male mice reared under ELA conditions were found to spend less time huddling with their cagemate than Control males (Welch’s t-test: t_(8)_ = 2.612, p=0.030; Figure 4A). In addition to reduced huddling duration, there was a trend for ELA males to be slower to initiate huddling (t_(14)_ = 2.072, p=0.057; Figure S1A). However, ELA and Control males spent a similar percent of time engaged in active interactions, and combined, there was no difference in the overall percent of time spent interacting (Figure 4B-C). To determine if there were differences between the various types of active interactions across the two male conditions, various types of active interactions were analyzed separately. No differences were found for the percent of time spent allogrooming (Figure 4D), or the number of investigative anogenital sniffs (Figure 4E), nor the latency to initiate either behavior (Figure S1B-C). However, there was a trend towards increased fighting behavior in ELA males (Welch’s t-test: t_(12)_ = 1.98, p=0.072; Figure 4F). Overall, these findings indicate that, following an acute stress challenge to the cagemate, ELA males engaged in less huddling behavior and instead engaged in more agonistic behaviors that resembled fighting.

**Figure 4.**
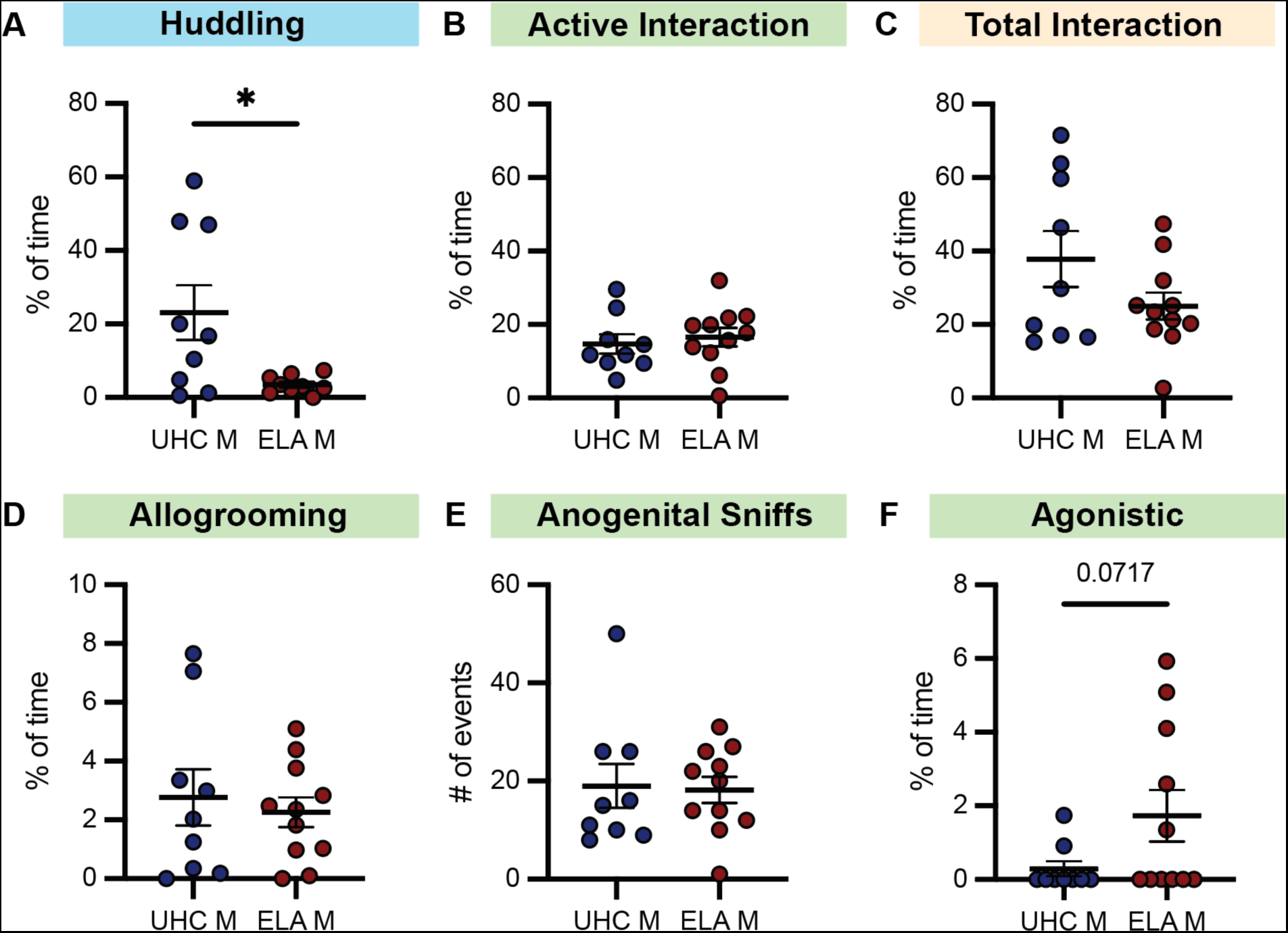
Social interaction between Males following acute stress exposure. Social interactions between the Shocked and Unshocked male cagemate were scored for 20 minutes following the footshock. The percent of time engaged in various behaviors were scored, including the amount of time spent engaged in: passive huddling (A), active interaction (B) and the combined social interaction time (C). Specific types of active interactions were also scored. The percent of time spent allogrooming (D), the number of anogenital sniffs (E) and the percent of time fighting (F) were quantified. A) ELA males spent significantly less time huddling than controls. B-C) The amount of time engaged in active interaction and total interaction did not differ based on rearing condition. Time allogrooming (D) and sniffing (E) also did not differ, however agonistic behavior tended to increase in ELA males (F). (*p<0.05). p-values for trends (0.10>p>0.05) are shown as values.

In contrast to males, Control and ELA females displayed a similar percent of time engaging in passive huddling behavior (n.s., p>0.05; Figure 5A). Interestingly however, ELA-reared females were slower to initiate huddling (t_(8)_ = 2.418, p=0.041; Figure S1D), similar to males. There was no difference in the percent of time spent actively interacting, nor in the total percent of interaction across the scoring session (Figure 5B-C). For the various active interactions, there was a decrease in allogrooming behavior in ELA females and an increase in the number of investigative anogenital sniffs (Allogrooming: t_(14)_ = 2.288, p=0.038; Sniffs: t_(13)_ = 2.301, p=0.039; Figure 5D-E). Mirroring the effects on duration, ELA females were also slower to initiate allogrooming behavior (t_(8)_ = 2.47, p=0.057; Figure S1E) and were faster to engage in anogenital sniffing (t_(10)_ = 1.90, p=0.0845; Figure S1F), though these findings did not reach significance. No fighting behavior was observed in any of the female cages, regardless of rearing condition (Figure 5F). Overall, these findings indicate that, following an acute stress challenge to one cagemate, ELA females engaged in less active prosocial behaviors such as allogrooming and instead engaged in more investigative behaviors like anogenital sniffing.

**Figure 5.**
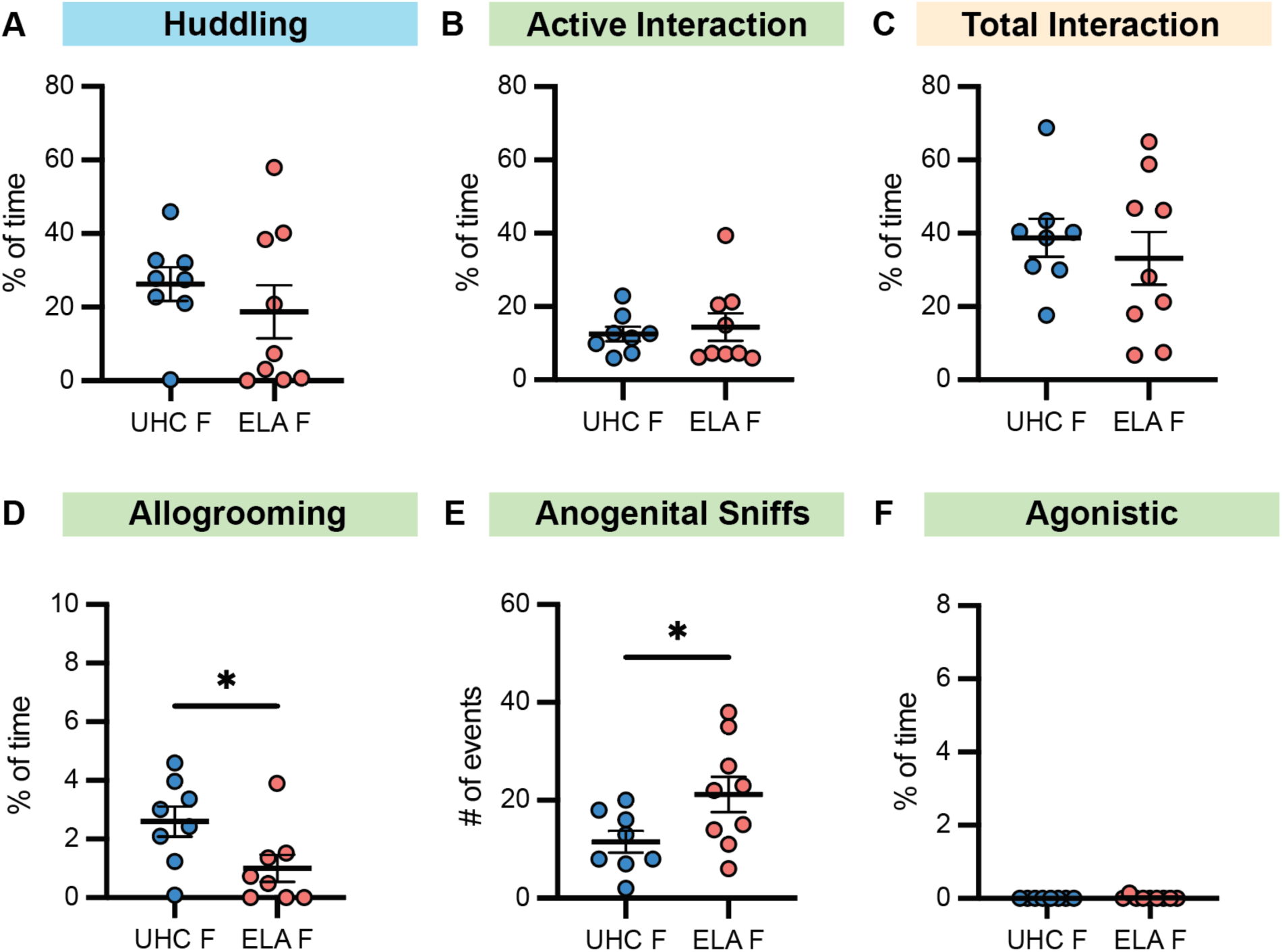
Social interaction between Females following acute stress exposure. Social interactions between the Shocked and Unshocked female cagemate were scored for 20 minutes following the footshock. The percent of time engaged in various behaviors were scored, including the amount of time spent engaged in: passive huddling (A), active interaction (B) and the combined social interaction time (C). Specific types of active interactions were also scored. The percent of time spent allogrooming (D), the number of anogenital sniffs (E) and the percent of time fighting (F) were quantified. The amount of time engaged in huddling (A) active interaction (B) and total interaction did not differ between ELA and control females. However, ELA females spent less time allogrooming (D) and had an increased number of anogenital sniffs (E). In both female conditions, agonistic behaviors were not observed (F). (*p<0.05).

## 4. Discussion

How an individual responds to an acute stressor can be highly variable (Sapolsky, 2015, 1994). Early life experience and biological sex likely contribute to such individual differences. Stress can also transfer across individuals and drive changes in social behavior, and social signals can serve to dampen or heighten stress responding (Burkett et al., 2016; Hall and Romeo, 2014; Romeo et al., 2006; Sterley et al., 2018). In the current study, we tested the effects of ELA and sex on the physiological stress response (measured by circulating CORT) following direct footshock or co-housing with a recently shocked cagemate, and examined social interactions during this experience. We found that ELA-rearing and sex differentially impacted circulating CORT levels. In males, both control and ELA-reared Shocked and Unshocked mice showed similar elevations in CORT at 20 minutes and a similar recovery at 90 minutes, indicating that a transfer of distress occurred, but was not impacted by ELA. In females, all groups again showed a significant rise in CORT at 20 minutes, however, Unshocked ELA females showed a higher CORT response compared to Unshocked controls, implying an increased transfer of stress as a result of ELA. In addition, ELA-reared females, both Shocked and Unshocked, maintained a heightened CORT response 90-minutes post stress, whereas Control females had returned to baseline levels. Differences in social behavior following footshock stress were also observed. ELA male mice displayed significantly less huddling and instead tended to display increased fighting behavior, though this did not reach statistical significance. ELA females engaged in less active prosocial behaviors such as allogrooming and instead engaged in more investigative behaviors like anogenital sniffing. Together, these findings suggest that a history of ELA alters the CORT response to a footshock stressor in a sex dependent manner, with female ELA-reared mice showing poorer regulation of the stress response. ELA rearing also altered the response to a distressed cagemate, exacerbating the physiological transfer of distress in female mice and reducing affiliative and prosocial behaviors in male and female mice compared to their respective controls.

### 4.1 ELA and the HPA axis

Various forms of ELA, including abuse and neglect, can have lasting effects on the HPA axis (reviewed in Fogelman and Canli, 2018; Koss and Gunnar, 2018). Adult humans who have experienced ELA generally show reduced cortisol levels in response to psychosocial stressors (Bunea et al., 2017; Carpenter et al., 2011; Lovallo, 2013), with increased ELA severity associated with greater blunting of CORT levels (Zhang et al., 2019). However, findings have been mixed (Fogelman and Canli, 2018), with potential moderating effects of variables such as stress experienced in adulthood (Goldman-Mellor et al., 2012) and co-occurrence of psychiatric disorders (Bremner et al., 2003; Heim et al., 2010, 2001, 2000). In addition, factors such as sex and the type of adversity experienced may differentially impact the HPA axis (Bourke et al., 2012; Neigh et al., 2009).

Rodent models of ELA have also produced mixed results regarding ELA’s impact on HPA axis functioning. For instance, studies using maternal separation (MS) have largely shown little to no change in adult stress responses (Parfitt et al., 2007). In contrast, rodents reared under the limited bedding and nesting (LBN) model, used here, have been reported to demonstrate measurable elevations in basal CORT (Bath et al., 2016; Molet et al., 2014; Moussaoui et al., 2017; Raineki et al., 2010; Rice et al., 2008). These elevations in basal levels are detectable around the time of the cessation of LBN manipulations. In response to stress challenges early in life however, both blunted (McLaughlin et al., 2016) and elevated (Gilles et al., 1996) CORT levels have been reported. Less work has examined how LBN affects stress responses later in life (post-weaning), though there is some evidence that LBN does not alter hormonal stress responses in adolescent (Molet et al., 2016b) or adult (Brunson et al., 2005; Eck et al., 2020) male rats. In the current study, we found that LBN-rearing did not impact baseline CORT levels in adult males or females. In addition, we found that male mice displayed similar CORT responses as their control counterparts following a stress challenge, matching prior literature.

Relatively few studies have tested the lasting effects of LBN on endocrine activity in female animals. A recent study by Eck and colleagues found no differences in CORT responses in LBN-reared female rats compared to controls following a 1-hour restraint stress, either immediately following the stress exposure or following a 30-minute recovery period (Eck et al., 2020). Consistent with these findings, here, we observed no effects of LBN on peak CORT responses in female mice at 20-minutes post stress. However, in contrast, we found that LBN-reared females failed to recover to baseline CORT levels. It is possible that differences in species (rats vs. mice) and the use of different stressors and sampling methods could explain these discrepancies. Additionally, in the study by Eck et al (2020), CORT collection only continued for 30 minutes after stress termination, and animals across all conditions never returned to baseline CORT levels. Thus, it is possible that LBN effects might have been observed if the time course of CORT measurement had been extended beyond the 30-minute period sampled. Furthermore, in the current study, mice interacted with their Unshocked cagemate following stress, perhaps contributing to the observed effects on the stress response. Regardless, our finding that LBN-reared females did not fully recover to baseline CORT levels 90 minutes after stress exposure has important implications. A sustained stress response, even after cessation of the stressor, could lead to altered reactivity to future stressors and impairments in fear extinction, among other proximate effects (Gump and Matthews, 1999; Maren and Holmes, 2016; Romero et al., 2009). A prolonged stress response, especially if repeated, can also have detrimental effects on health, leading to heightened allostatic load and increased risk for immune system dysfunction, cardiovascular disease, and mood disorders, among other conditions (De Kloet et al., 2005; McEwen, 2003, p. 200, 1998).

### 4.2 ELA and Emotional Contagion

Most studies examining how ELA alters the stress axis have tested stressors on isolated individuals. However, both humans and rodents exist in a social environment and social context can influence stress in myriad ways (Beery and Kaufer, 2015). For example, peers can help reduce or buffer another’s stress response, and stress can also transfer to an individual that was not the direct target of the stressor (Burkett et al., 2016; Hall and Romeo, 2014). The current study is the first, to our knowledge, to examine adult stress response in ELA-reared mice within the social context of a cagemate. In all conditions, Unshocked mice exposed to a Shocked cagemate displayed increased CORT levels, demonstrating a transfer of stress. In addition, CORT levels of the Shocked cagemate were positively correlated with that of their Unshocked partner in all conditions except, surprisingly, Control females, indicating that the magnitude of the stress response in the Shocked conspecific largely influenced their Unshocked cagemate; this is consistent with prior literature in rodents (Li et al., 2019). Overall, this transfer of distress, measured here by elevated and similarly matched CORT levels in the Unshocked cagemate, is indicative of emotional contagion. Emotional contagion refers to when witnessing an emotion in a conspecific elicits a similar emotion in the observer (Keysers et al., 2022), and is often studied in rodents by assessing whether an individual will demonstrate fear or pain-related behaviors upon observing a conspecific in distress, receiving a shock, or in pain respectively (Jeon et al., 2010; Keysers et al., 2022; Lu et al., 2018; Smith et al., 2021). Here, though the Unshocked mouse did not directly witness their cagemate receive footshocks, they interacted with the stressed animal upon their reunion, and the subsequent physiological stress response elicited in the Unshocked cagemate is indicative of stress contagion. This finding alone is important, as many studies use shocks as a means of testing learning and memory and return animals to group-housed conditions, potentially leading to stress contagion.

By and large, little is known about how ELA may influence emotional contagion. In the current study, LBN and Control-reared adult males displayed similar levels of emotional contagion (as indexed by transfer of the CORT response). This is in line with recent work by Laviola and colleagues (2021), which utilized a mouse model of ELA by providing dams with CORT in the drinking water for 1 week following parturition. Male animals reared by dams receiving CORT, or by control dams, were tested in a social contagion of fear paradigm in adulthood and no significant differences in paw licking (a measure of emotional contagion) were observed (Laviola et al., 2021). Here, we report for the first time that ELA led to both an increased transfer of distress in adult female mice, and an inability to regulate that response, as indexed by persistent elevations in circulating CORT levels. This sex-selective increase in CORT in Unshocked ELA females relative to Controls indicates that females, but not males, exposed to LBN may have heightened emotional contagion. Though others have not directly studied emotional contagion in female rodents reared under LBN conditions, one report by Lukkes and colleagues (2017) tested behavioral responses in MS and control-reared adolescent rats after witnessing a peer receive a series of footshocks. Interestingly, MS-reared female rats who had witnessed their peer receive a shock were faster to escape in a later active avoidance task, demonstrating reduced depressive-like behavior (Lukkes et al., 2017). Whether or not this behavior might be attributed to differences in stress contagion cannot be assumed and is an open area for future study. In general, how ELA affects emotional processing and emotional contagion remains an underexplored and potentially fruitful area of study. In future work, it would be important to characterize how ELA alters various types of emotional contagion, including social transmission of stress, fear and pain, as the neural circuits implicated in each differ (Keysers et al., 2022; Walsh et al., 2023).

### 4.3 The impact of ELA on Social Interactions

Experiencing ELA can profoundly alter social interactions later in life. Across species, ELA generally leads to reduced sociality and diminished social interactions (Bolton et al., 2018; Molet et al., 2016a; Patterson et al., 2022; Perry et al., 2019; Shin et al., 2018; Yazgan et al., 2021). However, the type of ELA experienced, and the age and sex of the individual may yield different outcomes in social behavior (Bondar et al., 2018; Holland et al., 2014). The current study expands on this body of literature and is the first to assess how ELA might change social interactions with a distressed peer.

Rodents typically respond to distressed conspecifics by displaying increased affiliative and prosocial behaviors such as huddling (side-by-side contact) and allogrooming (Burkett et al., 2016; Carneiro De Oliveira et al., 2022; Du et al., 2020; Lim and Hong, 2023; Lu et al., 2018; Matsumoto et al., 2021; Wu et al., 2021). Here, we found that mice reared under ELA conditions showed reduced affiliative and prosocial behaviors when interacting with a stressed peer. Specifically, ELA males huddled less, while ELA females reduced their allogrooming behavior. It is possible that these changes in social behavior are due to reduced social bonds between ELA cagemates, making ELA mice less likely to huddle or allogroom regardless of the circumstance. Future studies would be needed to investigate this possibility more rigorously, as prior work showing reduced social interaction in LBN-reared mice has focused on adolescents and has not analyzed home cage behavior in adult cagemates (Bolton et al., 2018; Molet et al., 2016a). Additionally, the diminished affiliative behaviors of ELA female and male mice cannot be attributed to a lack of physiological response to stress, as all groups showed a robust stress response in the current study. Indeed, the transfer of distress from the Shocked mouse to their Unshocked cagemate may have contributed to observed changes in social interaction. For example, it is possible that the heightened stress response in Unshocked ELA females prompted a reduction in allogrooming, though huddling and other active investigation measures remained unchanged. Relatedly, both huddling and allogrooming have been proposed as coping behaviors that can reduce stress responses (Beery and Kaufer, 2015; Morrison, 2016; Zeng et al., 2021). Thus, the reduction in allogrooming observed in ELA females could have contributed to the observed prolonged CORT response in the distressed (Shocked) cagemate. Importantly, no fighting was observed in the female groups, and thus, the sustained elevation of CORT in ELA females is likely not due to increased aggression. It is also possible that huddling and allogrooming might not be associated with the CORT responses observed here. In particular, the function of allogrooming in mice remains debated, as it can also be observed in dominant animals towards their subordinate cagemate (Lee et al., 2019).. Regardless, as cages were measured as single units for social behavior, the current study was underpowered to directly test how CORT responses related to social behavior, and thus, each of these proposed possibilities remains unexplored and will be an important area for future work.

In addition to the observed changes in affiliative behaviors, LBN-reared males also tended to display heightened agonistic, aggressive behaviors towards their distressed cagemate. Curiously, despite the higher levels of fighting and the reduced huddling in ELA males, all males showed similar CORT dynamics, indicating that these social behaviors likely did not influence CORT levels. This increase in agonistic behavior is also in contrast with prior work showing that ELA in the form of maternal separation can decrease agonistic behavior in adult male mice (Veenema, 2009; Veenema et al., 2007). These differences may stem from the model of ELA used and the familiarity of the animals; prior work has tested aggression towards an unfamiliar intruder, whereas here we tested aggression towards a cagemate. In addition, here agonistic wrestling and fighting was only observed in a subset of ELA males. Future work will need to increase sample size to better understand individual variability in agonistic behaviors (including the degree and type of agonism), with the goal of identifying characteristics of ELA mice that are prone to fighting or not and the conditions under which agonism is increased or decreased. Lastly, though agonistic behavior was not observed in any female mice in the current study, we did observe mounting behavior in a subset of control and ELA females, a behavior that has been linked to social hierarchies in females (Williamson et al., 2019). Thus, for both typically reared animals and those reared under ELA conditions, it will be important to continue to characterize how exposure to a distressed peer alters social interactions.

Altogether, our findings suggest that ELA may have lasting effects on the development of social behavior, reducing consolation-related behaviors towards a distressed peer. These findings have important implications for understanding the long-term consequences of ELA on social behavior and highlight the need for further research, both in animal models and in human populations.

### 4.4 Possible neural mechanisms underlying differences in the CORT response

The different temporal components of the stress response (baseline levels, stress reactivity and recovery) provide different windows of insight into the functioning of the stress axis, with distinct physiological processes and neural mechanisms underlying each phase (Herman et al., 2020; McEwen, 1998). Here, we observed sex-specific changes in CORT responses in ELA-reared adult mice, with footshock-exposed ELA females showing a prolonged CORT response. This impaired termination of the stress response could be due to alterations in glucocorticoid receptor (GR) and/or mineralocorticoid receptor (MR) density in the hippocampus, as they play an important role in HPA axis recovery (Gjerstad et al., 2018; Sapolsky et al., 1984). Hippocampal GR expression is suppressed both in individuals with a history of childhood abuse (McGowan et al., 2009) and in animal models of ELA (Avishai-Eliner et al., 1999; Bath et al., 2016; Enthoven et al., 2010; Francis et al., 1999; Schmidt et al., 2002; Van Oers et al., 1998; van Oers et al., 1997). However, none have tested whether female animals reared under LBN conditions have long-lasting changes in GR/MR expression; such information would aid in the interpretation of the CORT findings observed here. In addition, here, Unshocked ELA females had a heightened transfer of distress, with increased CORT after both 20 and 90 minutes of interaction with the distressed animal, relative to controls. This heightened social transmission of physiological stress in ELA females indicates there may be ELA specific changes in neural circuits associated with emotional contagion (such as in the anterior cingulate cortex and insular cortex) and/or in circuits associated with the social transmission of stress (such as in corticotrophin-releasing hormone (CRH) neurons in the paraventricular nucleus (PVN) (Carrillo et al., 2019; Paradiso et al., 2021; Rogers-Carter et al., 2018; Sterley et al., 2018). Whether and how LBN-rearing alters such neural circuitry, and whether it is associated with the observed changes in CORT observed here, remain critical areas of future study.

### 4.5 Possible mechanisms underlying differences in social behavior

In the current experiment, ELA altered social interactions between Shocked and Unshocked cagemates. Specifically, ELA females showed increased anogenital sniffing of their distressed cagemate. In general, investigative, exploratory behaviors are often observed when rodents are exposed to a distressed individual (Burkett et al., 2016; Knapska et al., 2006), and anogenital sniffing in particular may be associated with pheromones released from the anogenital region that allow unstressed partners to detect stress signals (Inagaki et al., 2009; Kiyokawa et al., 2004a; Kiyokawa, 2017; Sterley et al., 2018). Whether footshock exposed ELA females release more pheromones than their control counterparts, potentially driving this increase in anogenital sniffing remains an open question. Alternatively, this increased investigative behavior may be driven by a heightened exploratory drive in the Unshocked cagemate. Recent work has also shown that anogenital sniffing is required for the social transmission of stress; exposure to a swab from the anogenital region of a stressed mouse was sufficient to induce stress transfer (Lee et al., 2021; Sterley et al., 2018). Thus, the increased social investigation in ELA females observed here may underlie the heightened stress response observed in the Unshocked female cagemates, an association that will need to be tested in future work.

In addition to changes in anogenital investigation, we also found that ELA inhibited affiliative behaviors towards distressed peers, reducing allogrooming in ELA-reared females and reducing huddling in ELA-reared males relative to controls. Both proximity seeking and allogrooming of a distressed conspecific are associated with activity in brain regions involved in emotional contagion, including the amygdala and insula, among others (Lim and Hong, 2023; Matsumoto et al., 2021; Rogers-Carter et al., 2018; Wu et al., 2021). Oxytocin receptors are critical for partner directed allogrooming behavior, especially in brain regions such as the anterior cingulate cortex (ACC), insular cortex, lateral septum and medial amygdala (Burkett et al., 2016; Matsumoto et al., 2021) and LBN was found to alter oxytocin and oxytocin receptor positive cells in an age and sex specific manner (Lapp et al., 2020). Thus, in future work it will be important to explore whether ELA drives changes in neural activity in these regions, which could potentially explain the observed reductions in affiliative behaviors.

### 4.6 Limitations and Future Directions

In the current study, we observed effects of ELA on both CORT responses and social behavior of adult mice exposed to a shocked cagemate. However, there are several limitations that should be considered. Most significantly, we were underpowered to meaningfully explore correlations between physiological stress responses and social behavior. As mentioned previously, there are several possible means by which ELA induced disruptions in the HPA axis may contribute to altered social development and social behaviors. Thus, future studies should directly assess whether observed changes in CORT are correlated with alterations in social behavior towards a distressed conspecific. In addition, here we assessed CORT responses 20 minutes and 90 minutes after reunion with a shocked cagemate, chosen based on time course studies in our lab and others (Shanks et al., 1990). Future work should increase the number of timepoints analyzed to better understand the duration of the observed prolonged CORT response in ELA females. Furthermore, here, social behavior of shocked and unshocked mice were analyzed as a unit, and thus, it remains unclear which stress condition drove changes in social behavior. Behavior of individual animals should be tracked in future work.

Here, we show that CORT responses of shocked mice robustly transfer to an unshocked cagemate, indicative of stress contagion. However, pairing a stressed mouse with a non-stressed cagemate could also lead to social buffering, defined as an attenuation of the stress response when in the presence of another (Liu and Yuan, 2016; Morrison, 2016). Indeed, adverse childhood experiences may influence the capacity for others to provide stress-buffering benefits. Both human and animal studies have indicated that early adversity can diminish maternal buffering of stress, such that maternal presence fails to robustly reduce CORT responses to a stressor (Gunnar et al., 2015; Opendak et al., 2017; Sanchez et al., 2015). Less is known regarding how ELA might impact social buffering and the associated neural systems in adulthood. One recent study found evidence supporting LBN-induced alterations in social buffering in adult, female rats (Lapp et al., 2020), and studies have shown that both huddling and allogrooming behavior can help to buffer stress responses (Burkett et al., 2016; Hennessy et al., 2009; Kiyokawa et al., 2004b; Kiyokawa and Hennessy, 2018; Morrison, 2016). Thus, one hypothesis generated from the current study is that the reduced huddling and allogrooming behavior seen here in LBN-reared male and female mice, respectively, might lead to impaired buffering and downstream effects on stress-sensitivity.

### 4.7 Implications for Humans

The current study adds to a growing body of literature exploring the impact of ELA on HPA axis functioning and social interaction. While ELA did not affect peak CORT levels, both Shocked and Unshocked female ELA mice failed to return to baseline levels 90 minutes after stress exposure, suggesting that ELA-reared females exhibit difficulty in regulating the stress response. These findings have implications for understanding the long-lasting consequences of ELA on the HPA axis in humans. Impaired HPA negative feedback leads to protracted HPA axis responses and has been linked with psychiatric disorders such as major depression (Burke et al., 2005; Juruena et al., 2021; Pariante, 2009; Pariante and Miller, 2001). Future work will thus be needed to understand whether ELA-induced changes in HPA axis functioning are linked with augmented risk for the development of psychiatric disorders following ELA (Carr et al., 2013; Heim and Nemeroff, 2001; Targum and Nemeroff, 2019; Ventriglio et al., 2015; Watson et al., 2007).

Here, we also observed an increased transfer of physiological stress in ELA females, which may be indicative of heightened stress reactivity when faced with a distressed conspecific. This finding is consistent with evidence suggesting that children exposed to ELA are more sensitive to negative emotions and struggle with emotional regulation (Marusak et al., 2015; Tottenham et al., 2010). A transfer of distress is also indicative of emotional contagion, which is often considered to be a building block of empathic behavior, and our findings here could imply that ELA more broadly leads to enhanced emotional contagion. However, it is important to note that a transfer of distress does *not* imply empathy (Keysers et al., 2022). Indeed, though research on how ELA might alter emotional contagion is limited, there is some evidence that human adults with a history of ELA show *decreased* compassion and emotional empathy (Ardizzi et al., 2016; Fourie et al., 2019; Grimm et al., 2017; Luke and Banerjee, 2012; Parlar et al., 2014). Interestingly, one study found that children with a history of maltreatment displayed increased autonomic activity when viewing negative facial expressions, while simultaneously showing reduced empathic measures (Ardizzi et al., 2016). In line with this idea, here, despite evidence for a strengthened transfer of distress, we observed reductions in consolation-related behavior in ELA mice, with ELA-reared males huddling less and ELA-females allogrooming less than controls. These findings may indicate that ELA leads to a heightened reactivity to the stress of another, while simultaneously inhibiting affiliative social behaviors. Moreover, these reductions in social interaction are also consistent with human findings showing that individuals with a history of ELA have changes in social interactions, including increased antisocial behavior, poorer social relationships and smaller social network sizes (Ebbert et al., 2019; Ford et al., 2011; Kendall-Tackett, 2002; Larsen et al., 2011; Yazgan et al., 2021). However, whether ELA exposed individuals are more affected by the stress of their peers, and whether ELA leads to altered stress coping, remain open questions.

## 5. Conclusions

In sum, our results suggest that ELA may lead to long-lasting changes in stress reactivity and social behavior in adult mice. These findings have broader implications for understanding how ELA affects stress coping, physiological responding to stress, emotional contagion, and empathy-related social behaviors in humans. Further research in this area, in both humans and animal models, is greatly needed and would help shed light on the complex relationship between ELA and emotional development, with implications for understanding risk for psychiatric disorders.

## Acknowledgements

We would like to thank Dr. Giulia Zanni, Dr. Gordan Barr and Dr. Amiel Rosenkranz for their thoughtful comments on the manuscript.

## Author contributions

Conceptualization: J.M.B. & K.G.B. Methodology: J.M.B., C.D., R.R., K.G.B. Investigation: J.M.B., C.D., R.R., M.C., Z.C., K.D., S.N., D.O. Analysis: J.M.B. Visualizations & Writing: J.M.B. & K.G.B. Resources, funding and supervision: K.G.B.

## Data and materials availability

All analyzed data associated with this study are available in the main text or in the supplementary materials. Raw data will be made available upon request.

