## Supplemental Material for "Early life adversity reduces affiliative behavior towards a distressed cagemate and leads to sex-specific alterations in corticosterone responses"

| Dam ID | Rearing Condition | Litter # | Total # Pups in litter | Total # Pups used | Baseline | 20 min | 90 min |
|---|---|---|---|---|---|---|---|
| 211_C | Control | 5 | 9 | 9 | 1 (1F) | 6 (4F, 2M) | 2 (2F) |
| 213_C | Control | 3 | 10 | 8 | 2 (1F, 1M) | 6 (2F, 4M) | - |
| 246_C | Control | 4 | 9 | 7 | 1 (1F) | 4 (4F) | 2 (2F) |
| 248_C | Control | 3 | 5 | 2 | - | 2 (2M) | - |
| | Control | 6 | 1 | 1 | - | - | 1 (1M) |
| 263_C | Control | 1 | 5 | 3 | 1 (1F) | 2 (2F) | - |
| | Control | 2 | 6 | 5 | 1 (1M) | - | 4 (2F, 2M) |
| | Control | 5 | 5 | 5 | - | - | 5 (4F, 1M) |
| 266_C | Control | 1 | 9 | 7 | 1 (1F) | 6 (2F, 4M) | - |
| | Control | 4 | 9 | 9 | 1 (1M) | - | 8 (4F, 4M) |
| 267_C | Control | 1 | 8 | 8 | 2 (2M) | 6 (2F, 4M) | - |
| 283_C | Control | 1 | 8 | 4 | - | - | 4 (4M) |
| 288_C | Control | 1 | 10 | 6 | - | - | 6 (2F, 4M) |
| 216_E | LBN | 5 | 10 | 10 | 2 (2M) | 8 (4F, 4M) | - |
| 233_E | LBN | 3 | 9 | 4 | - | - | 4 (2F, 2M) |
| | LBN | 4 | 12 | 9 | 3 (2F, 1M) | 6 (4F, 2M) | - |
| 234_E | LBN | 5 | 7 | 6 | - | - | 6 (2F, 4M) |
| 238_E | LBN | 4 | 9 | 9 | 1 (1F) | 8 (4F, 4M) | - |
| 253_E | LBN | 2 | 4 | 2 | - | 2 (2M) | - |
| 255_E | LBN | 4 | 8 | 8 | 2 (1F, 1M) | - | 6 (2F, 4M) |
| 256_E | LBN | 2 | 8 | 8 | 2 (1F, 1M) | 6 (4F, 2M) | - |
| | LBN | 6 | 4 | 3 | 1 (1M) | - | 2 (2M) |
| 265_E | LBN | 2 | 11 | 4 | 2 (2F) | 2 (2M) | - |
| | LBN | 4 | 6 | 6 | 2 (1F, 1M) | - | 4 (2F, 2M) |
| 291_E | LBN | 1 | 9 | 9 | 1 (1F) | - | 8 (6F, 2M) |

**Supplemental Table S1. Dam and animal information across all conditions.**

Each unique dam and their rearing condition (control or LBN) is reported, as well as the litter number, the total number of pups in that litter and the number of pups used in the current experiment. For each of the three time conditions (baseline, 20 min or 90min) the number of total number of animals used is reported, along with sex in parentheses (M = male, F = female).

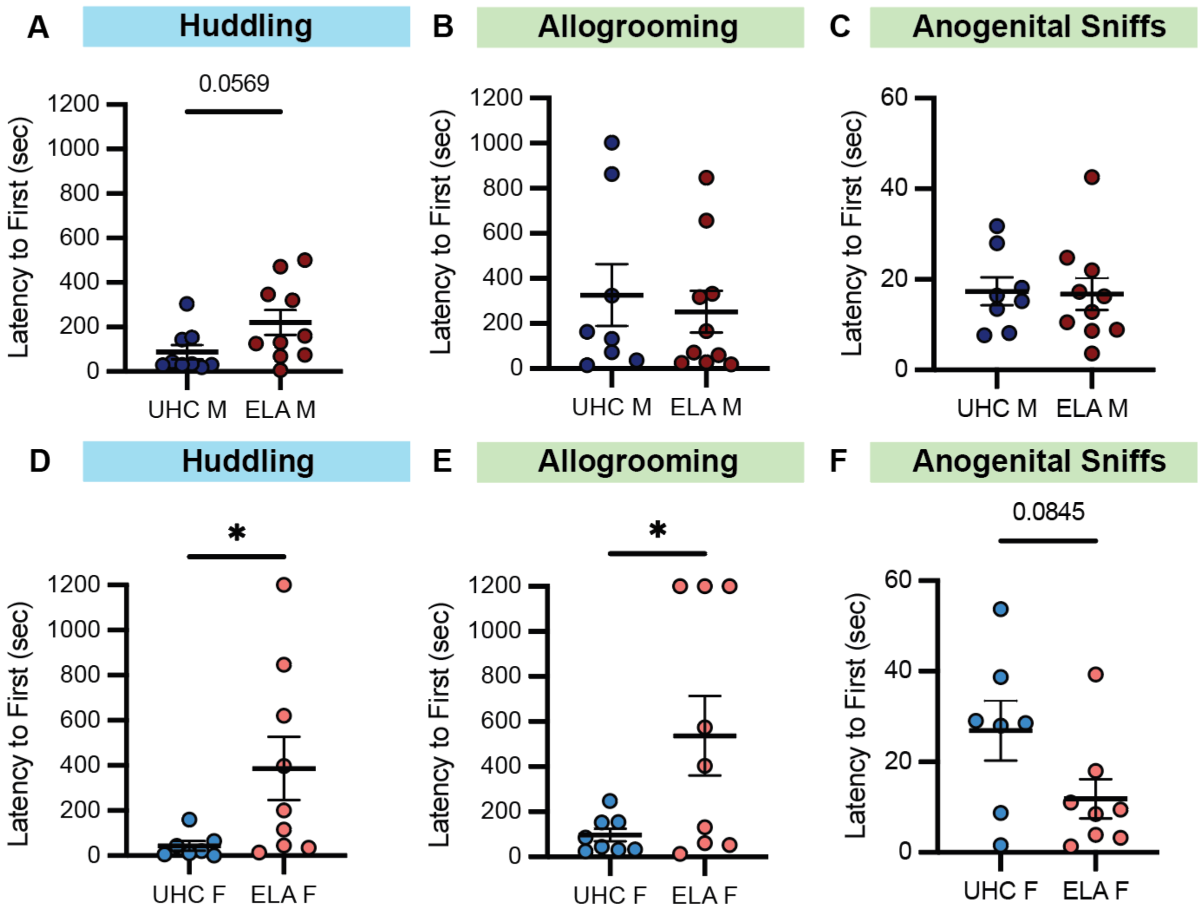


**Supplemental Figure S1. Latency to first social behavior.**

The latency to the first instance of each social behavior was quantified for both male (A-C) and female (D-F) mice. The first instance to huddle (A,D), allogroom (B,E), and sniff the anogenital region (C,F) are shown. A) ELA males showed a trend towards an increased latency to huddle. B-C) In males, there was no difference in latency across groups for allogrooming or anogenital sniffs. D) ELA females were significantly slower to initiate huddling. ELA females were also slower to allogroom (E) but trended towards being faster to sniff (F). (*$p<0.05$). p-values for trends ($0.10>p>0.05$) are shown as values.
